# Is an individual recognition immutable? Subordinate hermit crabs fight with a familiar but weaponless dominant

**DOI:** 10.1101/2025.05.14.654137

**Authors:** Chiaki I. Yasuda

## Abstract

Individual recognition can help subordinates avoid contests with lower success against familiar dominant opponents. This holds true in crustaceans, but if the dominants autotomize their weapon (chela/cheliped) before a second encounter, there is a decrease of potential fighting ability in the prior dominant and an increased chance of success in the prior subordinates than before. I examined whether familiar recognition is still effective even when subordinates encounter a familiar, but temporarily weaponless, dominant through two consecutive trials of male–male contests in the hermit crab *Pagurus middendorffii*. Subordinate *P. middendorffii* intruders avoid fights with a familiar dominant guarder. After establishing hierarchy and familiarity between subordinates and dominants possessing a weapon (major cheliped), the subordinates encountered the same dominants after experimentally induced autotomy of the weapon. Subordinate intruders actively fought with the familiar but weaponless guarders. This result might be explained by the subordinate’s updating information; that is, opponents were recognized the same but weaker than before. Information updating could be a useful new line of investigation into cognitive ability in invertebrates.

## Introduction

Dyadic contests cost energy and time for the contestants [1], so there is interest in how animals avoid contests with low chances of success [2–5]. When two contestants have already constructed a dominance hierarchy, individual recognition would be very useful to a subordinate for avoiding aggression against a familiar dominant in a second encounter [6,7]. According to Tibbetts and Dale [8], subordinates learn a recognition cue of a dominant associated with the dominance hierarchy. Individual recognition in this context is therefore assumed to the dominant’s fighting ability, and hence the hierarchy, at least until the next encounter.

However, this might not apply to crustaceans, although many species show individual recognition associated with a dominance hierarchy [9–12]. Members of this taxon use a cheliped as a weapon during contests. Although the presence and/or the size of the cheliped often overcomes a body size disadvantage [13–15], the weapon can be easily lost, even within a few minutes, by autotomy [16], unlike other weapons such as horns, antlers, or mandibles [17]. Therefore, if a dominant has autotomized its cheliped before a second encounter, its potential fighting ability should be much lower than at the prior interaction. This means a greater chance of success by a subordinate than before, leaving two options: avoiding a fight again based on the prior hierarchy, or initiating a fight regardless of the prior hierarchy. No studies deal with this issue as far as I know despite ubiquitous autotomy in crustaceans [18].

Male–male contests of *Pagurus* hermit crabs are an ideal system for examining the relationship between recognition and changing weapon status. *Pagurus* males show precopulatory mate guarding during the reproductive season, and a solitary intruder often initiates a fight to take over a guarder’s female [19]. During these contests, both contestants use a major cheliped as a weapon, and weaponless contestants have reduced fighting success [20– 22]. In *P. middendorffii*, male weapon size is a more reliable predictor of contest outcome than male body size [23], and subordinate intruders show individual recognition, with avoidance of a familiar dominant guarder during a second encounter, but not with an unfamiliar guarder [24]. However, autotomy of the weapon often occurs through predator avoidance when an individual is grasped by a predatory crab [25].

I here examine the behavioral response of a subordinate intruder against a familiar but weaponless dominant guarder in *P. middendorffii* to assess whether individual recognition by a subordinate intruder is still in effect even when a prior hierarchy are not guaranteed. To accomplish this, I conducted trials using two-sequential trials of male–male contests where the dominant in the first trial was experimentally induced to autotomize its weapon before the second trial.

## Materials and methods

### Experimental procedure

I collected precopulatory guarding pairs of *P. middendorffii* from the intertidal rocky shore at Kattoshi, southern Hokkaido, Japan (41°40′N, 140°36′E) in November 2016, the peak of the mating season at this site [26]. In the laboratory, if a male was intact and still guarding a female, I separately maintained the male and the female of each pair in a container (14.3 × 10.8 × 7.2 cm) or plastic cups (300 ml), respectively, to prevent copulation before the experiment. After the laboratory experiments, I measured the shield length (SL, calcified anterior portion of the cephalothorax, index of body size) of all individuals to the nearest 0.01 mm.

The experimental procedure is similar to that of Yasuda et al. [24] (see also the Appendix in the study for the descriptions of male–male contest in the *P. middendorffii*). To examine the effects of a weaponless familiar dominant guarder on the behavior of a subordinate intruder, I established two groups (autotomy, *n* = 40; control, *n* = 47). In both groups, I randomly assigned two pairs collected the same day as a set, and the smaller or larger male in each set was chosen as the focal intruder (SL = 3.37 ± 0.78 mm, mean ± SD) or the guarding male (SL = 4.72 ± 0.96 mm), respectively. In Trial 1, I placed a guarding pair in an arena (19.5 × 12.0 × 7.0 cm) with seawater (3 cm deep). After confirming guarding behavior, I introduced an intruder and recorded the outcome of the contest at 10 min from its initial movement. No intruders ended up guarding a contested female in Trial 1 (i.e., they all lost), therefore all intruders from Trial 1 were determined to be subordinate to the dominant guarders. After Trial 1, there was a one-hour interval until Trial 2.

During the interval, I induced guarders in the autotomy group to autotomize their major cheliped by grasping it with forceps without excessive force (i.e., not manual declawing; [27]). Since *P. middendorffii* autotomize own major cheliped when they are grasped by a predatory crab [25], this procedure was mimic this natural response and used in my previous study [28]. Autotomy occurred within five minutes. Precopulatory guarding in *Pagurus* is conducted by using a minor cheliped [29], so weaponless guarders could still possess a female as usual. I grasped the major cheliped of guarders in the control group by forceps and quickly released. No crabs in either group lost other appendages during the procedure. Trial 2 was then initiated with the same methods as in Trial 1, with the subordinate intruder from Trial 1 encountering the familiar dominant guarder either with or without a weapon.

I recorded all trials using a digital camera (DMC-LF1, Panasonic) and observed up to 10 min from the first movement of the intruder. If an intruder initiated grappling with a guarder (for details of this behavior, see [23]), I considered that the trial had escalated. If contestants did not show aggression for more than three minutes, I defined the fight as settled and recorded the escalation duration (sec) and the eventual outcome, based on which male ended up guarding the female. Cases where contestants were still grappling at the end of Trial 2 were defined as a draw because both males held the shell of the contested female.

### Data analyses

I used a model selection approach based on Akaike’s information criterion (AIC, [30]). I first examined the factors affecting contest escalation by using a generalized linear mixed model (GLMM) with a binomial error distribution. The response variable was whether intruders initiated grappling (yes = 1, no = 0, *n* = 87 × 2 = 174). The explanatory variables were (1) Group (autotomy = 1, control = 0), (2) Trial (Trial 1 or 2), and (3) Group × Trial interaction. To examine the effects of male fighting ability and resource value, I also treated (4) the difference in SL between intruder and guarder (DSL_I–G_) and (5) SL of the guarded female (SL_F_) as explanatory variables in the model. Intruder ID was treated as a random effect in the GLMM.

In escalated contests, I also used model selection based on the AIC for analysis of the contest duration for both trials (*n* = 64, Cox’s model with mixed effects) and the eventual outcomes after escalation in Trial 2 (*n* = 19, generalized linear model [GLM] with a binomial error distribution). Because all intruders lost in Trial 1, only the outcome in Trial 2 was used in the latter analysis. The response variable was the duration of a series of escalations (mixed Cox’s model, sec) or the outcome for intruders (GLM, win = 2, draw = 1, lose = 0). In the mixed Cox’s model, the explanatory variables and random effect were the same as in the GLMM (i.e., [1]–[5] and intruder ID). The explanatory variables in the GLM were (1) Group, (2) DSL_I–G_, and (3) SL_F_. All analyses were performed using the software R v.4.1.1 [31]; the R packages “glmmML” [32] and “coxme” [33] were used for GLMM and mixed Cox’s analyses. In Cox’s model, the proportional hazard assumption was satisfied for all explanatory variables (*P* > 0.23).

## Results

Table 1 summarizes the results of male–male contests. The model that best described whether intruders escalated a contest was the model with Group, Trial, Group × Trial, and DSL_I–G_ (*n* = 174, Table 2a). In Trial 2, subordinate intruders escalated contests more often in the autotomy group than in the control group (Table 1, Fig 1a), indicating that aggression by subordinates succeeded more against weaponless familiar guarders than against intact guarders.

**Table 1.** Summary of experimental groups and processes in male–male contests of *Pagurus middendorffii*.

| Group | N | Contest process |  |  |  |  |  |
| --- | --- | --- | --- | --- | --- | --- | --- |
|  |  | Escalation |  | Outcome of intruders in trial-2 |  |  |  |
|  |  | Trial-1 | Trial-2 | Win | Draw | Lose |  |
|  |  |  |  |  |  | Without escalation | With escalation |
| Autotomy | 40 | 21 | 16 | 4 | 8 | 24 | 4 |
| Control | 47 | 24 | 3 | 0 | 0 | 44 | 3 |

All intruders lost (i.e., they were not guarding a female) in Trial 1. After a one-hour interval, the subordinates encountered the familiar (same) dominant guarder in Trial 2. In the autotomy group, dominants were experimentally induced to autotomize its weapon (major cheliped) during the interval.

**Table 2.**
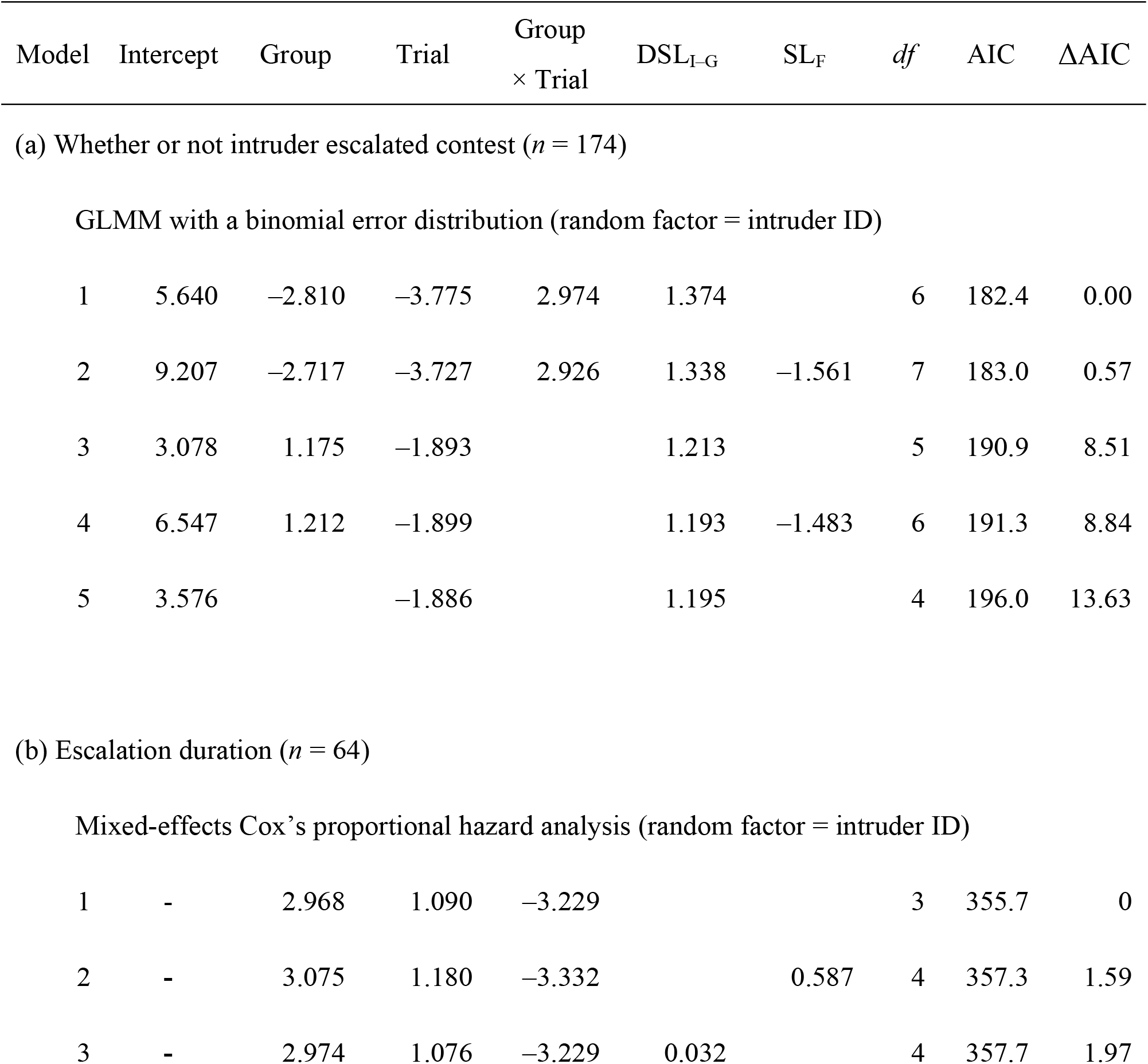
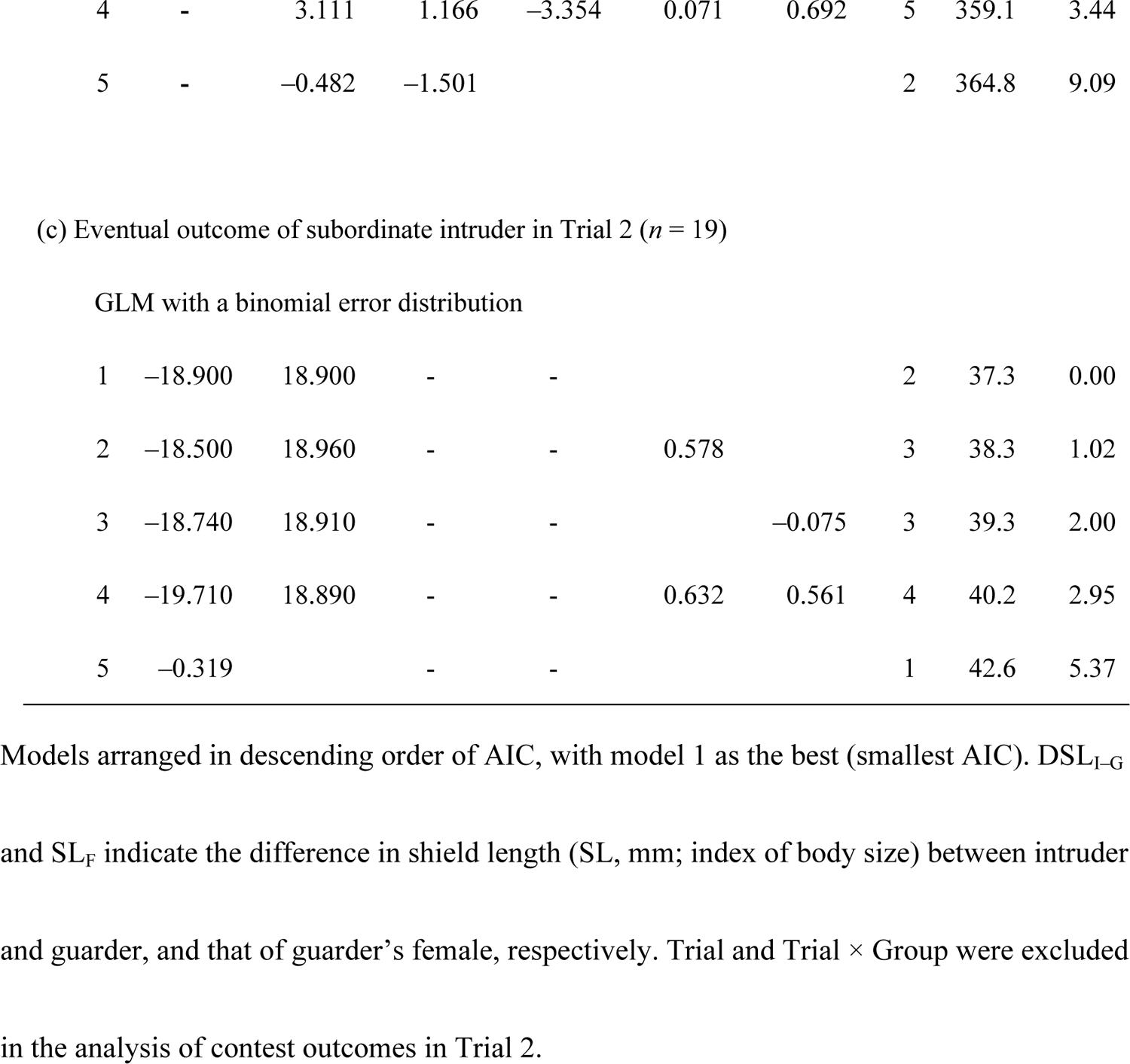
Results of the top five models based on Akaike’s information criterion (AIC).

**Fig 1.**
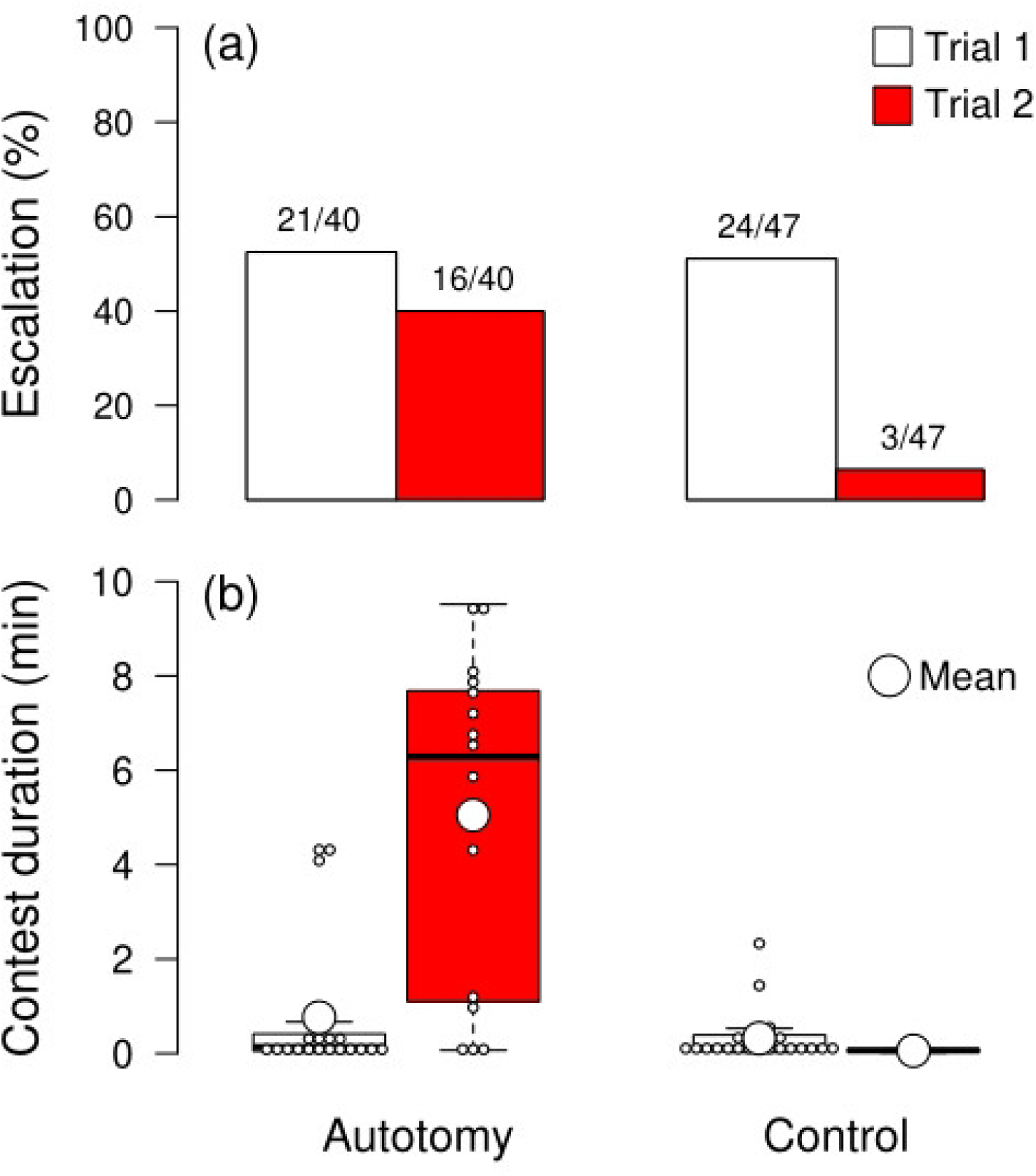
Differences in (a) intruder’s escalation frequency, and (b) escalation duration, between autotomy and control groups. Subordinate intruders (losers in Trial 1) encountered a familiar dominant guarder either with (control) or without (autotomy) its major cheliped in Trial 2. Numbers above bars in (a) indicate numbers of trials (escalated/total).

The duration of escalated contests was best described by the model with Group, Trial, and Group × Trial (*n* = 64, Table 2b). Contest duration in Trial 2 was much longer in the autotomy group than in the control group (Fig 1b), indicating that contests between subordinates and weaponless familiar dominants were not soon resolved.

The eventual outcomes after escalation in Trial 2 were best described by the model with Group as the explanatory variable (*n* = 19, Table 2c). Only three subordinates in the control group escalated the contest in Trial 2. After escalation, all three subordinates were defeated, whereas eight subordinates in the autotomy group (50%) reached a draw, and four (25%) succeeded in taking over a female from a weaponless dominant (Table 1).

## Discussion

The weapon status of dominants strongly affected the process and outcome of contests between familiar males in *P. middendorffii*: while subordinates avoided a familiar, intact dominant, they were likely to initiate, extend, and win a contest against a familiar but weaponless dominant. The few escalations and short contest duration in Trial 2 of the control group is consistent with our previous study [24] and with other decapods showing individual recognition [9–12,34,35], suggesting that the present study confirms individual recognition. The aggression in Trial 2 of the autotomy group, therefore, indicates that the effectiveness of familiar recognition was reduced in subordinates encountering a familiar but weaponless dominant.

The contest process in Trial 2 in the autotomy group might be related to both increasing subordinate aggression and decreasing defensive performance in weaponless guarders. Subordinate intruders might have assessed the lower fighting ability of guarders by visual and/or chemical cues indicating the absence of a weapon. Willingness to fight often increases with the chance of winning [36] or against a relatively weak opponent [37], including in hermit crabs [38], and my results are consistent with this general pattern. *Pagurus middendorffii* guarders may increase their aggression in the pre-fight phase after winning [24] (c.f. winner effect, [4]). However, a weapon is a crucial determinant of their fighting ability [23], as in other animals [17,36]. Its loss should therefore greatly decrease its actual fighting ability. Although I did not record guarder’s behavior, their weapon could contribute to prevent an opponent from approaching and grappling [28], so subordinates could more easily be able to grapple with the weaponless guarder. Taken together, the greater number of escalated contests in Trial 2 of the autotomy group suggests that decreased defensive ability from autotomy outweighs the winner effect. Once a contest escalated, such lower performance in weaponless guarders could cause longer escalation and lower success as in other *Pagurus* crabs [22,38].More research is necessary to reveal the relative importance of cues indicating a guarder’s weaponless status and decreased fighting ability in the decision-making of subordinates.

Recognition cues are still unclear in *P. middendorffii*, so it is premature to interpret the increased aggression in the autotomy group as loss of individual recognition. Chemical information is one of the most important recognition cues (chemical signatures, [34]), and familiar recognition often disappears when a familiar individual is presented with the chemical cue of an unfamiliar individual [39,40]. If *P. middendorffii* also uses chemical cues for recognition, autotomy of the major cheliped might be sufficient to change the chemical signature of familiar dominants.

Some animals, however, use visual cues for recognition [41,42], especially the “face” [43,44], even invertebrates [45]. Familiar recognition based on the face often ignores other body parts. For example, the crayfish *Cherax destructor* shows visual recognition based on a face but not a chela [46]. The social fish *Neolamprologus pulcher* use facial coloration as a signal for familiar-unfamiliar discrimination but not bodily traits [44]. Weaponless status in this study might not affect the dominant’s face. We therefore have two ideas to investigate in terms of what subordinates perceive in a weaponless guarder: (1) an unfamiliar weaponless guarder (i.e., prior information is discarded or not used), or (2) a familiar, but weaponless, dominant guarder (information about a specific dominant is updated). Knowing whether animals can recognize changes in and update information about a recognized individual will provide new insights for studies of cognitive ability across taxa.

## Acknowledgements

I thank Prof. Satoshi Wada for field sampling and advice, Ayumi Morita for field sampling, and Yuko Kanchiku for assistance with experiments.

## Supporting information

**S1 data. Yasuda CI_data.txt**. Data of this study.

**S1 code. Yasuda CI_code.R**. R code of this study.

## Ethics statement

This work did not require ethical approval from a human subject or animal welfare committee.

## Author contributions

C.I.Y.: conceptualization, data curation, formal analysis, funding acquisition, investigation, methodology, project administration, resources, visualization, writing—original draft, writing— review and editing.

